# Global chromatin reorganization and regulation of genes with specific evolutionary ages during differentiation and cancer

**DOI:** 10.1101/2023.10.30.564438

**Authors:** Flavien Raynal, Kaustav Sengupta, Dariusz Plewczynski, Benoît Aliaga, Vera Pancaldi

**Author notes:** Joint Authors (Benoît Aliaga, Vera Pancaldi).

## Abstract

Cancer cells are highly plastic, allowing them to adapt to changing conditions. Genes related to basic cellular processes evolved in ancient species, while more specialized genes appeared later with multicellularity (metazoan genes) or even after mammals evolved. Transcriptomic analyses have shown that ancient genes are up-regulated in cancer, while metazoan-origin genes are inactivated. Despite the importance of these observations, the underlying mechanisms remain unexplored. Here, we study local and global epigenomic mechanisms that may regulate genes from specific evolutionary periods. Using evolutionary gene age data, we characterize the epigenomic landscape, gene expression regulation, and chromatin organization in three cell types: human embryonic stem cells, normal B-cells, and primary cells from Chronic Lymphocytic Leukemia, a B-cell malignancy. We identify topological changes in chromatin organization during differentiation observing patterns in Polycomb repression and RNA Polymerase II pausing, which are reversed during oncogenesis. Beyond the non-random organization of genes and chromatin features in the 3D epigenome, we suggest that these patterns lead to preferential interactions among ancient, intermediate, and recent genes, mediated by RNA Polymerase II, Polycomb, and the lamina, respectively.

Our findings shed light on gene regulation according to evolutionary age and suggest this organization changes across differentiation and oncogenesis.

## Introduction

Despite recent progress, our understanding of oncogenesis remains incomplete. The loss of cellular identity is a complex and multi-step process, and is characterized by processes at different levels (genetic, epigenetic, etc.). In the last decades, two major theories about oncogenesis were proposed: the oncogenes paradigm, which views cancer primarily as a genetic disease, and epigenetic oncogenesis (Capp et al., under review). Evolutionary biologists also hypothesize that cancer is the result of a breakdown of multicellular cooperation (Athena Aktipis et al., 2015), consistent with the fact that most cancer genes date back to unicellular organisms and the transition to multicellularity and complex body plans (Domazet-Lošo and Tautz, 2010; Yin et al., 2019; Henikoff, 2023).

The approximately 20,000 human coding genes emerged at different evolutionary ages and evolved over time, notably with DNA mutations and duplication events. Gene and epigenome evolved differentially. Indeed, a subset of chromatin marks, like Polycomb deposited H3K27me3 and promoter-associated H3K4me3, were conserved more often than expected at orthologous regions with low substitution rates. Three chromatin marks (DNA methylation, H3K36me3, and H3K27ac) in mammal ESCs are independent of DNA sequence evolution (Endo et al., 2020). With the genomic phylostratigraphy method, genes can be dated by looking at their homologs across species (Domazet-Lošo et al., 2007). Other methods have been proposed in the literature to estimate evolutionary gene age, such as studying duplicated genes, as demonstrated by Juan *et al*. (Juan et al., 2013). In this paper the authors highlight a correspondence between gene age, replication timing, expression breadth across tissues and their propensity to accumulate copy number alterations. They suggest a scenario in which old, housekeeping genes are somehow protected from variations (both transcriptional and sequence-based), while younger genes are located in regions of high variability, promoting their evolvability and restricting the impacts of their evolution to specific tissues (Juan et al., 2013). The spatial organization of genes within the nucleus is also thought to influence the segregation of genes of different ages (Sima and Gilbert, 2014).

In 2017, Trigos *et al*. showed that the gene expression of evolutionary old genes increased in cancer, without explaining the underlying mechanism (Bussey et al., 2017; Trigos et al., 2019, 2017). Transcriptomics analyses of seven solid tumors showed a significant global increase in the expression of genes originating from unicellular organisms, while genes of metazoan origin were inactivated in cancer (Trigos et al., 2017). The widespread expression of unicellular genes in cancer appears to be a general pattern, but the specific up-regulation of these genes varies according to their function, indicating that this process is regulated rather than random (Benton et al., 2021; Trigos et al., 2017). More interestingly, cancer cells show expression of genes that are specific to embryogenesis and are not expressed in healthy adult tissues (Schoenhals et al., 2009).

These findings raised new interest in a theory that implicated the re-activation of unicellular programs in cancer, namely the atavistic theory from 1914 (Boveri, 2008), which saw the malignant tumor cell as a previously normal and “altruistic” tissue cell that changed into an “egotistical” mode with a loss of its specific functions. Recently, purely epigenomic mechanisms, namely the transient inactivation of Polycomb, were shown to be sufficient for the initiation of cancer in the fruit fly (Parreno et al., 2024), while embryonal programs are often implicated in dedifferentiation in cancer (Burdziak et al., 2023; Hunter et al., 2024; Patani et al., 2020) and 3D chromatin rearrangements are well-known hallmarks of cancer as they unlock phenotypic plasticity (Achinger-Kawecka and Clark, 2017; Hanahan, 2022).

Despite findings on the relationships between gene ages, expression levels and their regulation across oncogenesis, the underlying mechanisms are unclear. Both the local chromatin context and 3D organization of the genome within the nucleus have been shown to be cell type specific and to be altered during differentiation and oncogenesis, suggesting their potential role.

Recent work has shown that promoter DNA sequence characteristics, including general GC content, TATA-box motifs and number of TF binding motifs, can be predictive of gene expression variability across tissues, across individuals and across single cells in multiple species (Einarsson et al., 2022; Sigalova et al., 2020), suggesting that the local chromatin environment, as well as sequence characteristics can impact expression regulation and variability.

On the other hand, the global chromatin context of genes is likely to also play a role. The 2-meter-long chain of DNA folds into 3D space and coils inside the nucleus of each cell, giving rise to several hierarchical structures, which can be different in each cell but can also globally reflect cell type and cell state (Beagan and Phillips-Cremins, 2020; Misteli, 2020). Despite consistent progress in our understanding of chromatin organization, there are still open questions about how the characteristics of the 3D epigenome can be related to cellular processes and phenotypes (Pancaldi, 2021; Papadogkonas et al., 2022; Zheng and Xie, 2019). To investigate this, a useful framework involves representing chromatin as a network in which nodes are chromatin fragments and edges are experimentally detected spatial contacts between fragments (Pancaldi, 2023).

Here, we investigate potential local and global epigenomic mechanisms that could mediate the regulation of genes of specific evolutionary ages. We consider human embryonic stem cells, several differentiated immune cells and leukemic cells to study the gene evolutionary classes’ epigenomic characteristics, their spatial patterns inside the nucleus, and their regulation across differentiation and oncogenesis. Integrating chromatin 3D structure data with epigenomic features using network approaches, we propose that different mechanisms could be responsible for the physical organization of genes of specific evolutionary ages. Specifically, ancient genes of unicellular origins and with house-keeping functions are mostly regulated and clustered by RNA polymerase 2, intermediate metazoan genes, often involved in oncogenesis, are mostly coordinately repressed and clustered by Polycomb factors and, finally, novel mammalian specific genes, often found to be more peripheral in the nucleus, are bound and coordinately repressed by the lamina.

## Methods

### Gene Ages

We used the gene age dataset from Trigos *et al*., which estimated gene ages through phylostratigraphy (Trigos et al., 2017). Here, 19.404 genes are classified into 16 age classes, from unicellular genes to Homo sapiens specific ones, which are grouped into three main gene age categories: Unicellular Ancestor (UC, 7.397 genes), Early Metazoan (EM, 8.682 genes), and Mammal-Specific (MM, 3.325 genes).

### Expression and DNA methylation Variability

Expression and methylation variability in monocytes, neutrophils and T-cells were computed from 125 healthy individuals, using a method that accounted for confounding effects due to the correlation between mean and variability measurements (Alemu et al., 2014; Ecker et al., 2017). Fisher’s exact tests were performed using the geneOverlap (v1.32.0) R package (Li Shen, 2017).

### ChIP-Seq Processing

Histone modifications ChIP-Seq datasets in monocytes were downloaded from the ENCODE portal (https://www.encodeproject.org/) (Sloan et al., 2016) in BigWig format, with the ENCODE identifiers listed in Supplementary Table 1. Other ChIP-Seq datasets were downloaded in FASTQ raw format. Sequenced reads were mapped to the human reference genome (built hg19/GRCh37) using Burrows-Wheeler Aligner (bwa v0.7.17-r1188) (Li and Durbin, 2010). BAM files were obtained, sorted, and indexed by using Samtools utilities (v1.15.1) (Li et al., 2009). ChIP-Seq peak calling was performed with MACS2 (v.2.2.7.1) using the respective inputs (Zhang et al., 2008). H3K27me3 peaks were called with the --broad parameter on.

### ChIP-Seq Analysis

ChIP-Seq profiles were generated using the seqplots R package (v1.23.3) (Stempor and Ahringer, 2016). Polycomb target genes were defined as genes with strict TSS localization, according to EnsDb.Hsapiens.v75 Ensembl hg19 gene annotations, that overlap with an H3K27me3 peak. CpG localizations were analyzed using the ChIPSeeker R package (v1.32.1) (Yu et al., 2015).

### RNA-Seq Analysis

The raw read count matrix for hESC and hESC-derived cardiomyocytes was downloaded from GEO (accession number GSE69618) (Busser et al., 2015). Differentially expressed genes were found using the DESeq2 R package (v1.36.0), filtering for log2FC > 2 or <-2 and adjusted p-value < 0.1. Gene Ontology enrichment analysis was conducted using the limma (v3.52.1) (Ritchie et al., 2015) and clusterProfiler (v4.4.3) (Wu et al., 2021) R packages.

### PolII Pausing Index

We downloaded raw RNA Polymerase II ChIP-Seq FASTQ files for hESC (H9/WA09 cell line), B-cell (GM12878 cell line), and Lymphoma B-cell (LY1 cell line) and processed them as described above. PolII BigWig density files were generated using the bamCoverage tool from deepTools suite (v3.0.2) and normalized with the --normalizeUsing RPKM flag. To compute PolII Pausing Indexes, we defined the promoter region for each gene as TSS-30bp to TSS+300bp and the gene body region as TSS+300bp to TES, based on EnsDb.Hsapiens.v75 gene annotation. PolII scores for overlapping bins within each region were averaged separately. The Pausing Index was calculated as the ratio of PolII density in the promoter region to the density in the gene body region. Outliers in each cell’s Pausing Index distribution were detected and removed using the InterQuartile Range (IQR) method. Genes with a sum of PolII Pausing Indexes across the three cell types lower than 0.2 were considered unexpressed and were excluded. Additionally, genes shorter than 1kb, according to EnsDb.Hsapiens.v75 gene annotation, were discarded. To facilitate comparison across cells, each distribution was normalized to fall between 0 and 1.

### LAD-associated genes

Nuclear lamina-interacting genes were defined as genes with at least part of the gene body overlapping with LADs, according to EnsDb.Hsapiens.v75 Ensembl hg19 gene annotations and published ‘constitutive’ and ‘facultative’ LADs, either together or separately (Kind et al., 2015).

### PCHi-C chromatin networks and ChAs calculations

PChi-C datasets used with GEO accession numbers are summarized in Supplementary Table S1. Considering significant interactions with CHiCAGO scores>5, we constructed networks by considering DNA restriction fragments as nodes and interaction between fragments as edges. These networks were processed using the igraph R package (v1.3.2) (Csárdi et al., 2023). Chromatin Assortativity was calculated using the ChAseR R package (v0.0.0.9) (https://bitbucket.org/eraineri/ChAseR/) (Madrid-Mencía et al., 2020). Gene age ChAs was computed using age classes in the categorical ChAs mode. For z-score calculations, we performed 100 randomizations preserving genomic distances (dist.match = TRUE). Chromatin networks were visualized using Cytoscape (v3.8.0) and Gephi (v0.10.1).

### ΔChAs calculation

We developed the ΔChAs method to identify chromatin features associated with the 3D clustering of specific gene groups (e.g., age categories) by determining whether these features significantly contribute to clustering a gene group together (ΔChAs > 0) or not (ΔChAs < 0). Within each category, two gene subgroups are created based on the presence or absence of a chromatin feature. ChAs z-scores for both subgroups are then calculated. Finally, ΔChAs is determined as the difference between the ChAs z-score of genes associated with the specific feature and the ChAs z-score of other genes.

## Results

### Gene expression and expression variability are strongly associated to evolutionary ages

To explore the interplay between genes’ evolutionary history, their regulation, and nuclear organization, we considered a dataset established through phylostratigraphy that assigns ages to genes based on their homologous relationships across species. The dataset comprises 19,404 human genes categorized into 16 evolutionary ages, which are further grouped into three principal age classes: unicellular (UC, 7.397 genes), Early Metazoan (EM, 8.682 genes), and Mammal-specific (MM, 3.325 genes) (**Supplementary Fig. 1A**) (Trigos et al., 2017). Throughout the manuscript, we will primarily focus on the three major gene age groups for clarity and relevance in specific analyses.

We performed a functional enrichment analysis of genes within each category (**Supplementary Fig. 1B**), confirming that UC genes are enriched in fundamental cellular processes (catabolism, RNA splicing, histone modification) (Trigos et al., 2017; Yin et al., 2019). This contrasts with later phylostrata genes, which are associated with more complex cellular functions (sensory perception, keratinization). Additionally, we explored the enrichment of housekeeping genes across all age classes (**Supplementary Fig. 1C**), observing a notable proportion of UC genes classified as housekeeping compared to EM and MM genes. (18% of UC genes against 6% of EM genes and 1% of MM genes), while the EM class contains the majority of proto-oncogenes within the leukemic context (Jang et al., 2015), with some others being in the UC class. (**Supplementary Fig. 1D**).

Healthy cells display abundant variability and plasticity in their phenotype, partly based on changes in the epigenome (Ecker et al., 2018, 2017), while expression variability can be related to cancer aggressiveness (Ecker et al., 2015). We therefore proceeded to investigate the relationship between gene age, mean expression and expression variability. Across 125 healthy individuals (Chen et al., 2016) (**Fig. 1A**, **Supplementary Fig. 1E**), our analysis revealed an inverse correlation between the 16 gene ages and average expression levels across these cell types (Spearman test, Rho = −0.95, p-value < 2.2e-16 in monocytes), indicating that older genes exhibit higher average expression levels, consistent with their involvement in fundamental cellular processes.

**Figure 1:**
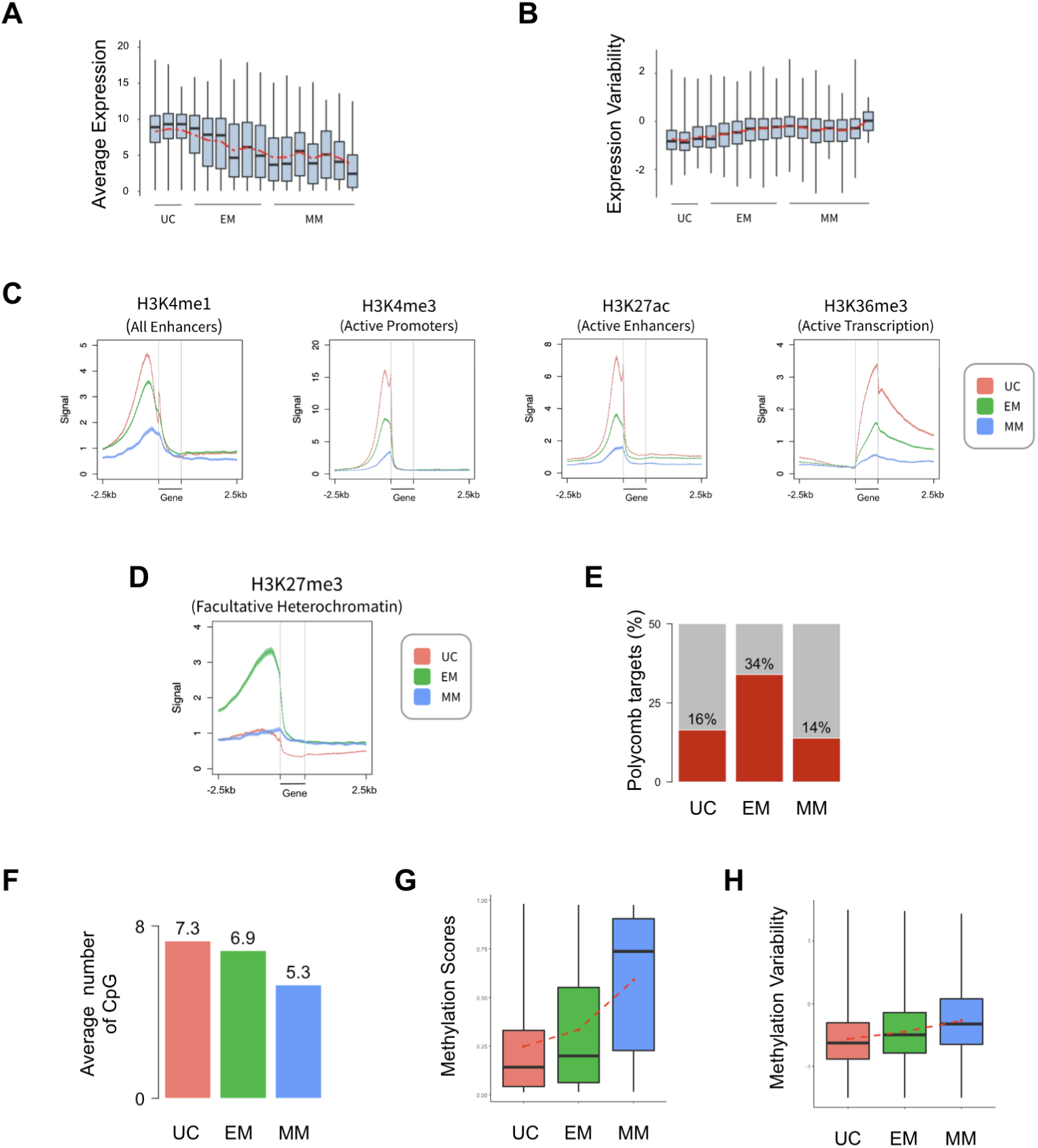
Genes of different evolutionary age classes display specific genomic and epigenomic features. **A**) Boxplot of average inter-individual gene expression levels in monocytes across 16 gene age classes (Spearman Rho = −0.95, *p*-value < 2.2e-16). **B**) Boxplot of inter-individual expression variability in monocytes across 16 gene age classes (Spearman Rho = 0.77, *p*-value = 0.00074). **C**) Average epigenetic profiles around the 3 main gene age classes (Red Unicellular genes, Green Early metazoan genes, Blue Mammal Specific genes) in monocytes. **D**) Average H3K27me3 profile around the 3 main gene age classes (Red Unicellular genes, Green Early metazoan genes, Blue Mammal Specific genes) in monocytes. **E**) Percentages of H3K27me3 repressed gene promoters in UC, EM and MM ages in monocytes. **F**) Average number of potential CpG islands by gene across the 3 main age categories in monocytes. **G**) Average Methylation scores computed by Chen *et al*. in monocytes across the 3 main age categories. Methylation scores were assigned to the 440,905 detected methylated CpG islands and calculated as the ratio between the intensity of the methylated probe and the global intensity within each individual. **H**) Average DNA methylation variability across the 3 main age categories in monocytes.

In the following, we define *variability* as variation of values across instances (across individuals or across single cells) and plasticity as variation in response to changes in external conditions. Briefly, Expression Variability (EV) (Alemu et al., 2014; Ecker et al., 2017) estimates the excess or reduced variability compared to what would be expected for the corresponding mean value of the feature. Since understanding expression variability is important for comprehending the underlying gene regulation mechanisms (Ecker et al., 2018), we examined the relationship between EV and age across immune cell types. (**Fig. 1B**, **Supplementary Fig. 1F, Supplementary Fig. 1G**), observing that older genes have lower variability compared to newer genes (Spearman test, Rho = 0.77, p-value = 0.0007 in monocytes). UC genes are significantly enriched in low variability groups (groups 1 to 6) while EM genes are enriched in high variability groups (groups 8 to 10) for each of the three immune cell types considered. These results were validated when considering gene subsets with different variability but similar expression (**Supplementary Fig. 1H**).

To further validate our findings, we conducted additional analyses using data from Sigalova *et al*. (Sigalova et al., 2020), which provides insights into variability across isogenic Drosophila lines and human tissues. Sigalova *et al*. identified an association between gene promoter features (sequence-based and epigenomic) in defining the level of gene expression variation. Analyzing these features related to variability in combination with the previously introduced age classes, we found correlations between gene age and active TSS (Spearman Rho = −0.37, p-value < 2.2e-16), the number of TF binding motifs in gene promoters (Spearman Rho = −0.34, p-value < 2.2e-16), the number of gene exons (Spearman Rho = −0.27, p-value < 2.2e-16), promoter size (Spearman Rho = −0.13, p-value < 2.2e-16) and GC content (Spearman Rho = −0.12, p-value < 2.2e-16) (**Supplementary Fig. 1I**).

Given the strong correlations observed among evolutionary ages, gene expression levels, and expression variability, we asked whether genes of specific age classes exhibit distinct epigenomic landscapes, which might imply different regulatory mechanisms (**Fig. 1C**). In monocytes, we found histone modifications associated with gene activation (H3K4me1/3, H2K27ac, H3K36me3) to be highly enriched around UC genes, intermediate intensity around EM genes and the lowest intensity in MM genes. These histone modification density levels are consistent with increasing expression levels for older genes, suggesting that the local epigenetic environment of the genes is also closely related to their evolutionary ages.

### Polycomb proteins target mainly metazoan genes

We then analyzed histone modifications associated to transcriptional repression (**Fig. 1D**, **Supplementary Fig. 1J**). Interestingly, in monocytes the H3K27me3 repressive mark shows enrichment around intermediate EM genes, while both older UC and younger MM genes exhibit no enrichment, suggesting a specific regulatory role by Polycomb complexes for this gene age category. Conversely, H3K9me3 mark shows no enrichment for any gene age class. This observation led us to focus on genes targeted by Polycomb proteins. We assessed the proportion of genes from each age class enriched with H3K27me3 peaks on the promoter in monocytes (**Fig. 1E**). Results reveal a higher proportion of EM genes (34%) targeted by Polycomb complexes whereas UC and MM genes exhibit a lower proportion of targeted genes (16% and 14%, respectively).

Taken together, these findings suggest that Polycomb protein complexes predominantly regulate genes that emerged during the metazoan period. This aligns with the role of these genes in establishing body plans and their regulated suppression in early phases of development.

### CpG islands and DNA methylation variability are associated to evolutionary ages

Next, we directed our focus towards DNA methylation as a key epigenetic feature. The dataset provided by Chen *et al*. contains methylation scores for each of the 440,905 detected methylated CpG sites across the entire human genome in monocytes, gathered from 200 healthy individuals via Whole Genome Bisulfite Sequencing (Chen et al., 2016). We first investigated the genome regions associated with these methylated CpGs which showed that half of all studied CpG islands are localized around gene promoters (52.52% within 3kb of the TSS, 42% within 1kb) (**Supplementary Fig. 1K**) and explored the correlation between gene ages and DNA methylation patterns. Methylated CpG islands are thought to repress gene expression when found within promoters (Bird, 2002). Our analysis indicated a progressive decrease in the average number of CpG islands within gene promoter regions across evolutionary gene age classes (**Fig. 1F**). We then analyzed methylation scores computed by Chen *et al*. in monocyte gene promoters classified according to their respective ages. Results show a gradual increase in methylation levels for newer evolutionary ages, suggesting a preferential regulatory role of DNA methylation for MM genes (**Fig. 1G**). Interestingly, inter-individual DNA methylation variability exhibited a roughly linear increase from older to newer gene age classes (**Fig. 1H**), suggesting a more stable DNA methylation profile for older genes and a greater variability around newer genes. This pattern may reflect an evolutionary process in which genes transition over time from being silenced in highly methylated regions for younger genes to more optimized and straightforward activation programs in older genes.

### Most genes deregulated through cell differentiation appeared recently in evolution

The above observations led us to focus on several gene expression regulatory mechanisms that could elucidate the organization and maintenance of age-related gene expression patterns. We investigated whether these patterns remain stable across global phenotypic shifts during cell differentiation and oncogenesis.

We studied three distinct and extensively studied cell states for comparison: human embryonic stem cells (hESCs, H9), representing an undifferentiated state, differentiated cells (CD19+ primary B-cells, monocytes, neutrophils, T-cells and cardiomyocytes), and B-cells from Chronic Lymphocytic Leukemia patients (B-CLL), as a representation of a cancerous leukemic cell state (Busser et al., 2015; Ecker et al., 2017).

We first asked whether genes of specific evolutionary ages are the most regulated during cell differentiation and oncogenesis. We analyzed a gene expression dataset from hESC differentiating into cardiomyocytes (Choy et al., 2018) and computed proportions of up- and down-regulated genes within each evolutionary age class observing a predominance of up-regulated genes across cell differentiation (from 5.382 differentially expressed genes, 89% are found up-regulated), which is consistent with acquisition of a lineage specific gene expression profile (**Fig. 2A**, **Supplementary Fig. 2A, Supplementary Fig. 2B**). Interestingly, this proportion of up-regulated genes rises across evolutionary age classes (14% of UC genes, 32% of EM genes and 59% of MM genes). Conversely, a minority of genes exhibited down-regulation during cardiac differentiation, with the number of down-regulated genes being even lower for younger gene age classes (5% of UC genes, 3% of EM genes and 2% of MM genes).

**Figure 2.**
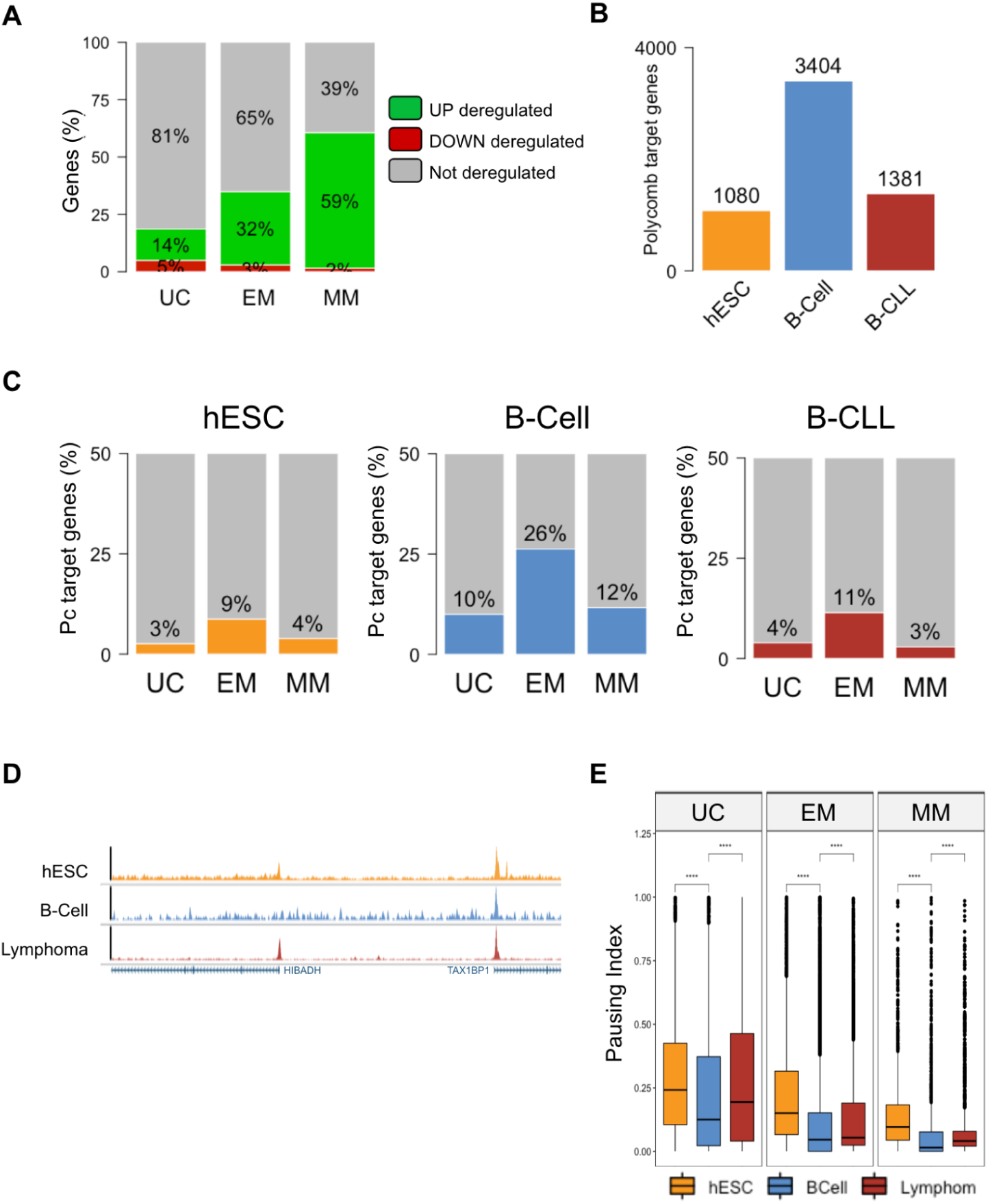
Gene expression regulation across different evolutionary ages is altered in differentiation and cancer. **A**) Proportion of down (red), up (green) and no (grey) deregulated genes in Cardiomyocyte compared to hESC for the 3 main gene ages. **B**) Number of Polycomb target genes in hESC (yellow), B-cell (blue) and CLL (red). **C**) Percentage of Polycomb target genes for the 3 main gene age categories in hESC (left), B-cell (middle) and CLL (right) cell states. **D**) Browser snapshots of PolII tracks in hESC (yellow), B-cell (blue) and CLL (red). **E**) PolII Pausing Index distributions in hESC (yellow), B-cell (blue) and CLL (red) for each of the 3 main age groups. Statistical comparisons between groups were done by the Wilcoxon-Mann-Whitney test. n.s = not significant, **** = p<0.0001.

### Gene expression regulation of genes from different evolutionary ages is altered in differentiation and cancer

In order to study how gene expression regulation is affected during cell differentiation and oncogenesis, we first focused on Polycomb protein complexes. We assessed the presence of the H3K27me3 mark in hESC, B-cell, and CLL as a proxy for Polycomb binding, which is known to repress developmentally regulated genes and influence expression noise levels (Kar et al., 2017; Kim and Kingston, 2022).

The total count of Polycomb-targeted genes across the genome was computed for each cell state, revealing an increasing number of targeted genes during cell differentiation and a sharp decline through oncogenesis (**Fig. 2B**). Building upon our earlier findings indicating that early metazoan genes (EM) were prominently regulated by Polycomb in monocyte (c.f. **Fig. 1D**, **Fig. 1E**), we observed a similar trend in stem cells, B-cells, and CLL cancer cells (**Fig. 2C**, **Supplementary Fig. 2C**), with the most notable impact of gene age class in B-cells compared to stem or cancer cells.

We then investigated gene expression regulation by RNA polymerase II (PolII) binding, another factor influencing transcription levels and variability, by analyzing Pausing RNA Polymerase II. This mechanism allows cells to modulate and synchronize transcription and can be quantified using the Pausing Index (PI) calculated as the ratio of PolII signal density around gene TSS to the PolII signal density along the gene body (Adelman and Lis, 2012).

PolII Pausing Indices were then computed for each gene in hESCs, B-cells, and B-CLL (**Fig. 2D**), and distributions were plotted for genes grouped by age class in each cell state, revealing a declining global distribution across evolutionary classes (**Fig. 2E**) and indicating that older genes are subject to tighter regulation by the PolII pausing mechanism. Interestingly, we observed significant decreases in global PolII pausing distributions during cell differentiation and significant increases during oncogenesis for every gene age class.

In summary, our findings indicate that the most recently evolved genes exhibit elevated expression levels during cell differentiation, while the oldest genes are most expressed during oncogenesis. EM genes are the primary targets of Polycomb complexes across all our cell states. Notably, the percentage of genes targeted by Polycomb seems to increase in differentiated cells and subsequently decrease in cancer. Additionally, our analysis suggests that PolII pausing is more pronounced in older genes, potentially influencing their expression regulation. With an opposite trend compared to Polycomb activity, this mechanism tends to decrease during cell differentiation and increase during carcinogenesis.

### The nuclear periphery is enriched with newer genes

Gene regulation, especially through Polycomb binding, has been strongly associated with the 3D chromatin structure within the nucleus, which was shown to be deeply compromised in Polycomb units mutants (Schoenfelder et al., 2015). Chromatin organization is known to be dramatically re-organised both in cell differentiation through lineage definition (Freire-Pritchett et al., 2017), and in cancer (Deng et al., 2022; Wang et al., 2022). This prompted us to investigate whether gene evolutionary ages might also be spatially organized in the nucleus.

To explore the potential association between gene evolutionary ages and genome 3D structure, we investigated whether genes from specific age categories are more likely to be situated at the nucleus periphery. We obtained a dataset of computed Lamina Associated Domains (LADs) annotated by Kind et al. (Kind et al., 2015), across nine human cell lines, describing LADs as either facultative (cell type-specific) or constitutive (cell type-invariant). We considered genes contained in facultative and constitutive LADs either together or separately and examined their age. Interestingly, we observed a higher proportion of younger genes associated with LAD domains (15% and 13% of MM and EM age categories, respectively) compared to older genes (7% of UC genes), (**Fig. 3A**, **Supplementary Fig. 3A**). Our findings support that genes located at the nucleus periphery are predominantly of newer origin and are consistent with the lower expression levels observed for the youngest genes (c.f. **Fig. 1A**).

**Figure 3:**
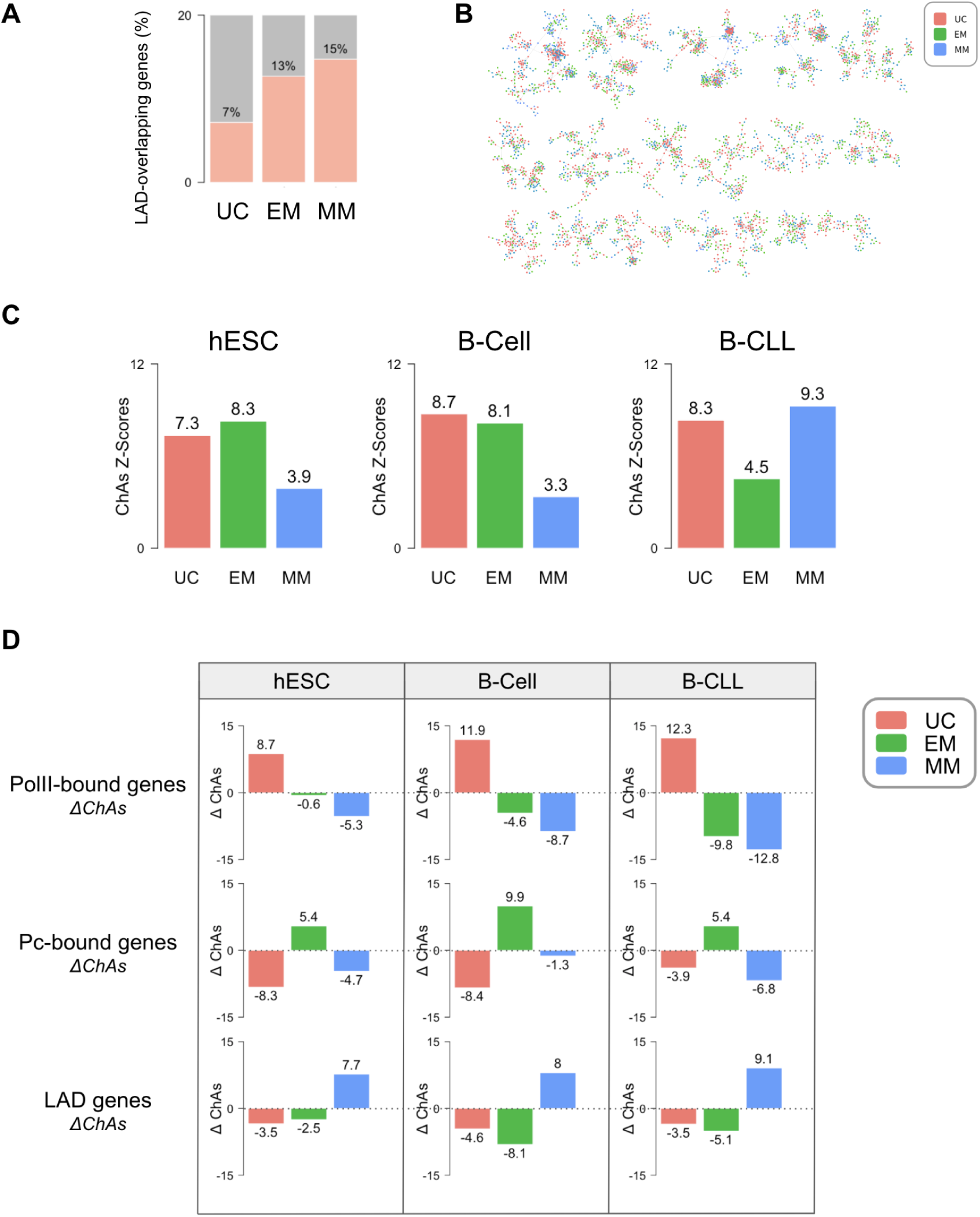
Spatial patterns of gene evolutionary age in 3D genome architecture. **A**) Overlap with Lamina Associated Domains (LADs) annotated in multiple cell lines (Kind et al., 2015) for genes belonging to the different age classes. **B**) Visualization of the 3D promoter-promoter interaction network in monocyte (Javierre *et al*. 2016) with node color representing gene age classes (red = UC, green = EM, blue = MM). **C**) Comparison of ChAs z-score of genes from different gene classes between hESCs, B-cells, and B-CLL. **D**) ΔChAs z-scores of different chromatin features: PolII (top), Polycomb (middle), and LAD (bottom) in hESC, B-Cell, and B-CLL.

### Genes establish preferential contacts in the 3D chromatin network according to their age

In prior studies, we and others have demonstrated that numerous epigenomic features including many transcription factor binding peaks, gene expression, DNA methylation, histone modifications (Madrid-Mencía et al., 2020), replication timing and replication origin activation (Jodkowska et al., 2022) display strong clustering in 3D space. We used the same network framework and chromatin assortativity (ChAs) to identify which features cluster more significantly on the chromatin network than would be expected at random considering solely linear proximity of the regions along the chromosome.

We exploited promoter-centered chromatin networks derived from PCHi-C datasets, where each node represents a DNA fragment and edges signify significant physical contacts (or proximity) between DNA fragments detected by the conformation capture protocol. We started by mapping gene ages on promoter only subnetworks in monocyte, neutrophil, and T-cell (**Fig. 3B**, **Supplementary Fig. 3B**) (Javierre et al., 2016) and observed that genes from specific age classes tend to group together, forming age-specific clusters. We next confirmed this tendency using ChAs computation with the three broad gene age classes interpreted as separate binary categorical values (**Supplementary Fig. 3C, Supplementary Fig. 3D**) (Madrid-Mencía et al., 2020). We observed ChAs z-scores for every age category being significantly above 2 (equivalent to p-value = 0.05), indicating that contacts are more frequent than would be expected given the genomic proximity of the regions and the number of promoters in each class. Interestingly, UC genes emerge as the most assortative gene class in all three cell types, consistently followed by EM genes. Conversely, the most recent gene age category displays lower assortativity across the three immune cell types.

These findings suggest that in fully differentiated cells the oldest unicellular genes exhibit stronger connectivity among themselves, forming prominent clusters in the chromatin network, closely trailed by EM genes, which also demonstrate preferential interactions. The newest genes have less pronounced preferential interactions, with ChAs z-score values just above significance.

Having observed specific interactions among genes of same evolutionary ages in monocyte, neutrophil, and T-cell, we next investigated the potential alteration of this age-related 3D conformational arrangement during cell differentiation and oncogenesis. We calculated ChAs z-scores on PCHi-C promoter subnetworks from hESC, normal B-cells, and cancerous B-CLL cells using the three primary age classes as binary categorical features (**Fig. 3C**). Across all cell states, we observed that each age category was significantly assortative. Notably, the oldest UC gene category consistently exhibited high assortativity values, indicating a persistent clustering tendency of these genes throughout cell differentiation and malignant transformation. Interestingly, EM genes display high ChAs values in hESC and B-Cells, and decline significantly during oncogenesis. Conversely, MM genes, which initially have the lowest values in both hESC and B-Cells, show a marked increase in clustering in CLL cancer cells. Overall, these findings indicate a substantial reorganization of interactions among both EM and MM gene groups during oncogenesis: strong clustering of EM genes is partially disrupted in cancer, while MM genes are reorganized into much stronger clusters.

So far, ChAs analysis has provided insights into chromatin features associated with 3D genome contacts, here highlighting that genes of specific ages cluster together. To further understand which specific proteins might be implicated in mediating this spatial organization, we performed a variation of ChAs analysis, denoted as deltaChAs (ΔChAs). Briefly, this method measures the difference in ChAs values obtained in subnetworks of chromatin fragments with or without a specific feature to determine whether it significantly contributes to clustering a gene group together (ΔChAs > 0) or not (ΔChAs < 0) (see Methods section for details).

We decided to test ΔChAs of several previously identified age-associated chromatin features in hESC, B-Cell, and B-CLL networks across each main age category (**Fig. 3D**). We observed positive ΔChAs for UC genes associated with PolII in all three cell types, suggesting a role of PolII in maintaining preferential contacts among UC genes in all cell states (c.f. **Fig. 3C**). The EM age class consistently exhibits positive ΔChAs z-scores for Polycomb targets compared to non-targets in all cell states, suggesting that the decrease of ChAs z-score for EM genes in B-CLL is primarily due to the disruption of preferential interactions among Polycomb-targeted EM genes which are preserved in hESC and B-cells. Lastly, despite their low abundance and weaker clustering in hESC and B-cells, we observed positive ΔChAs for MM genes associated with Lamina in all three cell states. This underscores the potential importance and specificity of LAD in the clustering of genes within the MM age category across different cell states, which is especially strong in CLL cells.

Positive ΔChAs values of EM genes for Polycomb observed in all cellular states suggest that the clustering of EM genes could be mediated by Polycomb across differentiation. Other factors might be important in mediating the clustering and repression of developmental genes, specifically in stem cells. Despite the limited number of ChIP-seq datasets publicly available, we were able to calculate ΔChAs in hESC and B-cells for several TFs, showing that several of them might contribute to maintaining the clustering of UC genes and EM genes, specifically TBP, KMT2A (hESC), and STAT3 (B-Cell) for UC genes, and CTCF (hESC) for EM genes (**Supplementary Fig. 3E)**.

In the context of 3D chromatin organization, our study reveals distinctive genome spatial patterns associated with gene evolutionary ages. Further analysis identified several chromatin features responsible for the 3D clustering of each specific evolutionary age. We observed that RNA Polymerase II facilitates preferential interactions among the oldest unicellular genes, aligning with their strong and stable expression, a trend consistent across various cell states. Lamina proteins could play a role in mediating interactions of mammalian-specific genes, showing moderate clustering in hESC and B-cell, and a notable increase in CLL. Polycomb proteins modulate the assortativity of early metazoan genes, a phenomenon particularly prominent in both undifferentiated and differentiated cells compared to cancerous states. This suggests a significant reorganization of early metazoan gene interactions driven by Polycomb activity alteration during oncogenesis.

## Discussion

### Investigating the link between gene age, regulation, and chromatin 3D organization

With the purpose of investigating potential mechanisms that relate genes’ evolutionary origins to their regulation and spatial organization we interrogated several datasets. Our results underscore relationships between evolutionary gene ages and several expression and epigenomic features, as well as with chromatin structure. We expand on previous observations linking gene ages to the TAD structure (Acemel and Lupiáñez, 2023; James et al., 2021; Sima and Gilbert, 2014) finding gene age to be globally assortative in chromatin networks, with particularly strong preferential clustering of genes of a particular age in specific cell states, and via regulation by potentially different molecular mechanisms.

Given the known earlier replication timing of older housekeeping genes (Juan et al., 2013) as well as evidence of 3D organization of replication timing domains (Pope et al., 2014), of replication origin activation (Jodkowska et al., 2022) and assortativity of several chromatin states (Pancaldi et al., 2016), these findings align with existing knowledge. Indeed, polycomb factors clearly associated with 3D structure have recently been implicated in origin activation (Prorok et al., 2023) and spatial organization of genes according to function, was suggested over a decade ago (Che et al., 2022; Cremer and Cremer, 2001; Sandhu et al., 2012; Takizawa et al., 2008) while the importance of genes’ radial positioning is also well known (Bouwman et al., 2022; Misteli, 2007) and altered in cancer (Meaburn et al., 2009). We here propose hypotheses on the factors establishing this organization and how it is modulated across differentiation and oncogenesis. Specifically, each gene age class exhibits distinct properties, which we will here summarize and consider in the context of relevant literature.

### Genes of unicellular origin: expression, regulation and alterations in cancer

The oldest genes, of unicellular origin, UC, display higher expression levels, lower inter-individual expression variability, and feature active chromatin marks. Exploration of a recent characterisation of expression variability across individuals and across tissues in combination with several epigenomic features suggests that old genes would display low variability thanks to having broad promoters and multiple regulatory motifs (Sigalova et al., 2020). It is well known that the oldest genes have functions related to basic processes in the cells, (housekeeping) (Ecker et al., 2018; Pancaldi et al., 2010; Yin et al., 2016). We find highest PolII Pausing Indexes, likely influencing their expression regulation. Interestingly, this mechanism tends to decrease during cell differentiation and increase during carcinogenesis. Previous studies demonstrated that the oldest genes, belonging to the UC age class, tend to exhibit higher gene expression levels across various solid tumor samples (Trigos et al., 2017) and were found to cluster on protein-protein interaction networks (Trigos et al., 2019). Specifically, our findings suggest that in all cell types and states the oldest unicellular genes exhibit more pronounced connectivity among themselves, forming prominent clusters in the chromatin network. Our results suggest that PolII binding is involved in clustering of UC genes while potentially maintaining coordination of their expression across differentiation and oncogenesis, in line with PolII roles in chromatin organization (Zhang et al., 2023, 2021). Despite our work focussing only on gene promoter regions, it has been recently shown that profiles of PolII intergenic binding are sufficient to cluster samples by cell line, tissue and cancer type (De Langen et al., 2023), suggesting that coordination of PolII bound genes could happen via complex networks of regulatory elements, as also suggested by analysis of PolII spatial patterns on mESC chromatin networks (Pancaldi et al., 2016). Moreover, it has been shown that establishment of replication timing in early development is impacted by PolII (Nakatani et al., 2024)

Overall our findings suggest that oncogenesis may reinstigate a balance characterized by high expression of old genes typical of undifferentiated states, which could lead to expression of ancestral and developmentally early programmes producing a shift towards unicellular phenotypes, in line with the atavistic theory (Lineweaver et al., 2021). These would be characterized by the Warburg effect and alterations in cell-cell communication as the tissue homeostasis is disrupted.

### Metazoan genes and Polycomb repression

Early Metazoan genes, EM, appeared with the evolution of multicellularity and complex body plans. They display intermediate expression and variability levels and enhanced Polycomb associated marks (H3K27me3). Polycomb protein complexes predominantly regulate genes that emerged during the metazoan period, aligning with the role of these genes in establishing body plans and their regulated suppression in early phases of development. Notably, the percentage of Polycomb targets seems to increase in differentiated cells and subsequently decrease in cancer, especially for EM genes. Those genes also demonstrate preferential interactions, but this pattern is lost in B-CLL cells, potentially in association with loss of cell identity. During development, Hox genes play a crucial role in forming the body plan (Mann and Glassford, 2024; Yamanaka et al., 2023) and rely on Polycomb group proteins for regulation (Gentile and Kmita, 2020; Kim and Kingston, 2022; Murphy and Boettiger, 2024). Moreover, numerous studies have also demonstrated that Hox genes are implicated in cancer (Brotto et al., 2020; Feng et al., 2021). Polycomb proteins could modulate the assortativity of EM genes, a phenomenon particularly prominent in both undifferentiated and differentiated cells compared to cancerous states. This suggests a significant reorganization of EM gene interactions driven by loss of Polycomb activity during oncogenesis, which was recently confirmed to produce cancers in drosophila in the absence of any genomic alteration (Parreno et al., 2024). The high value of ChAs z-score for EM genes in hESC aligns with earlier ChAs studies in mouse ESC (Pancaldi et al., 2016).

### Mammalian-specific genes: DNA methylation and chromatin reorganization

Lastly, the newest mammalian specific genes, MM, displayed higher levels of DNA methylation and methylation variability in their promoters, potentially explaining their overall lower expression level. The relationship between DNA methylation at promoters and gene expression remains poorly understood, with recent reports that sequence variants might underlie the observed negative correlation (Stefansson et al., 2024). However, the higher expression variability of new genes would be consistent with a less stable genome, accumulation of sequence variants, potentially through errors accumulated in late replication, and generally faster evolution (Juan et al., 2013). Recent peripheral enhancer regions characterized by H3K9me2 only might also play a role in regulation of MM genes (Smith et al., 2021).

Evidence from yeast also suggested a strong relation between variability of genes across single cells, across individuals, and in response to external changes (Barkai and Shilo, 2007) suggesting that genes related to immunity that have arisen more recently would display high variability in expression and faster genomic evolution (Hagai et al., 2018; Seufert et al., 2024).

During hESC differentiation into cardiomyocytes, genes in the MM class are predominantly up-regulated, reflecting the development of a lineage-specific gene expression profile. These findings may also be attributed to the increased variability of gene expression observed across pluripotent cells; low-level expression of MM genes might be due to different cells expressing different genes in hESC. During differentiation, lineage-specific genes become upregulated in all or most cells, resulting in strong upregulation at the population level.

Interestingly, we found genes located at the nucleus periphery enriched in the MM class. The importance of radial positioning of genes in relation to their transcriptional state has been known for close to two decades (Che et al., 2022; Takizawa et al., 2008). Given that genes interacting with nuclear lamina typically exhibit repressed expression (Reddy et al., 2008; Van Steensel and Belmont, 2017), this finding is consistent with the lower expression levels observed for the youngest genes (c.f. **Fig. 1A**). This association with the lamina might also in part explain the preferential interactions between MM genes, which is weak in stem and differentiated cells but becomes predominant in cancer, in which these genes are more strongly repressed as cell identity is lost. The effects of higher evolutionary plasticity of newer genes (Juan et al., 2013) might be controlled by their relegation to nuclear peripheral regions where they are only expressed in very specific conditions such as cancer cells. It is also possible that rewiring of chromatin in cancer alters the regulatory mode of specific gene classes, linking chromatin structure to expression noise levels via control of burst size and frequency (Seufert et al., 2024).

This work raises a series of hypotheses but we must mention several limitations of this study. First and foremost, we investigated published datasets integrating chromatin contact maps with epigenomic marks from different studies. These datasets may refer to cells in slightly different conditions and are likely influenced by biases specific to each type of experimental assay. Secondly, the chromatin assortativity measures depend, to a certain extent, on the size and characteristics of the contact networks being compared. To address this, we limited our comparisons to ChAs z-score values of gene age categories within a single chromatin network (e.g., hESC, B-cell, or B-CLL) rather than comparing across different networks. Thirdly, not all genes are present in the promoter networks, due to experimental limitations, potentially affecting our results. As a fourth point, when studying chromatin structure of cancer cells it is hard to guarantee that changes in chromatin conformation are not confounded with genome alterations. Additionally, we are here studying only the human genome 3D structure and chromatin organization principles have changed across evolution (Hehmeyer et al., 2023; Hoencamp et al., 2021), while single-cell specific chromatin organization cannot be captured by population-aggregated data, which only informs us on global trends and might be misleading (Messina et al., 2023). Finally, another limitation is due to the lack of datasets describing promoter-centered chromatin structure in primary solid cancer cells, which prompted us to look at a leukemic cell instead. Differences between B-cells and B-CLL cells might be less pervasive than those between healthy cells in a tissue and their oncogenic counterparts. If solid cancer phenotypes can be related to a loss of cellular identity due to alterations of the tissue context, in leukemia the tissue context is totally absent (other than in the bone marrow and in lymph nodes). This could explain why the re-expression of UC genes is not observed in this cancer type (data not shown).

## Conclusion

In conclusion, despite these limitations, the strong relationship we observed between gene age, expression variability, regulation and 3D localisation, combined with evidence that specific genes relocate within the nucleus during oncogenesis prompts us to propose that the reshaping of chromatin during oncogenesis may modulate the variability and plasticity of phenotypes. Further work will have to be devoted to determining whether changes in the regulation of specific genes or a global rearrangement of broad gene age categories in three-dimensional space contribute to the observed alterations in variability (across cells) and plasticity (over time) during differentiation and loss of cell identity, potentially sufficient to promote oncogenesis (Frank and Yanai, 2024).

## Data availability

No new data were produced for this study. The full list of previously published datasets used is provided in **Supplementary Table S1**. The script and processed datasets used for analysis and to generate main figures in this study are publicly available in Figshare at https://doi.org/10.6084/m9.figshare.26977654.

## Acknowledgments

The authors thank Michael N. Kammer, Daniel Rico, and David Juan for the critical reading of the manuscript.

## Author contributions

F.R, B.A. and V.P. designed the study. B.A. provided knowledge and background of the study. F.R. performed all bioinformatics analysis. K.S. and D.P. provided support with aspects of the network science expertise. F.R, B.A. and V.P. wrote the manuscript. V.P. supervised the project.

## Funding

This work was supported by the Fondation Toulouse Cancer Santé and the Pierre Fabre Research Institute, as part of the Chair of Bioinformatics in Oncology at the CRCT; Agence Nationale de la Recherche [ANR-23-CE12-0023, ANR-24-CE52-0683-02]; K.S. and D.P. were funded from Warsaw University of Technology under the Excellence Initiative: Research University (IDUB) programme and co-supported by the Polish National Science Centre [2020/37/B/NZ2/03757].

## Conflict of interest statement

The authors declare no conflicts of interests.

## Supplementary Figures

**Supplementary Fig 1. (related to Fig. 1).**
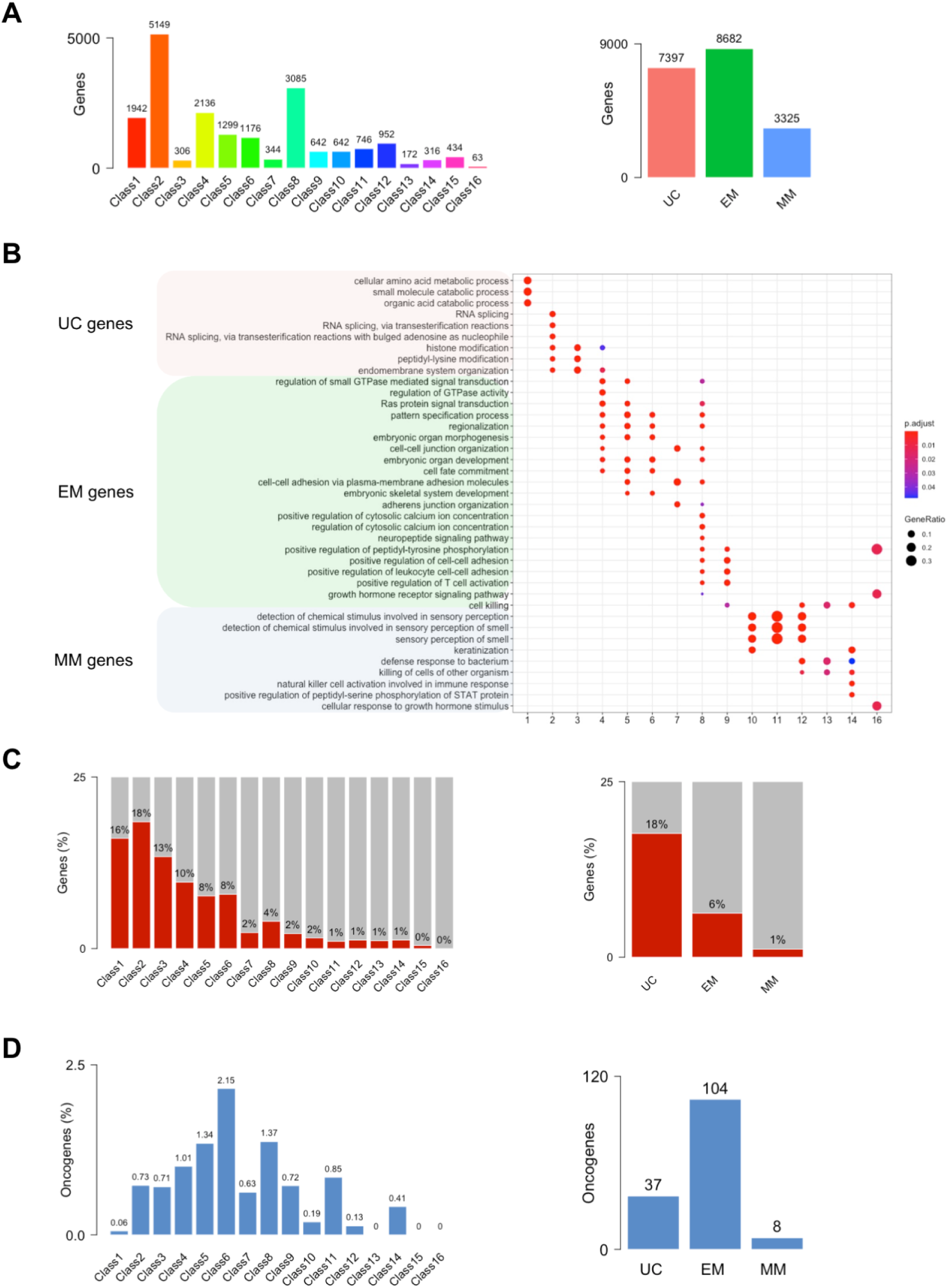
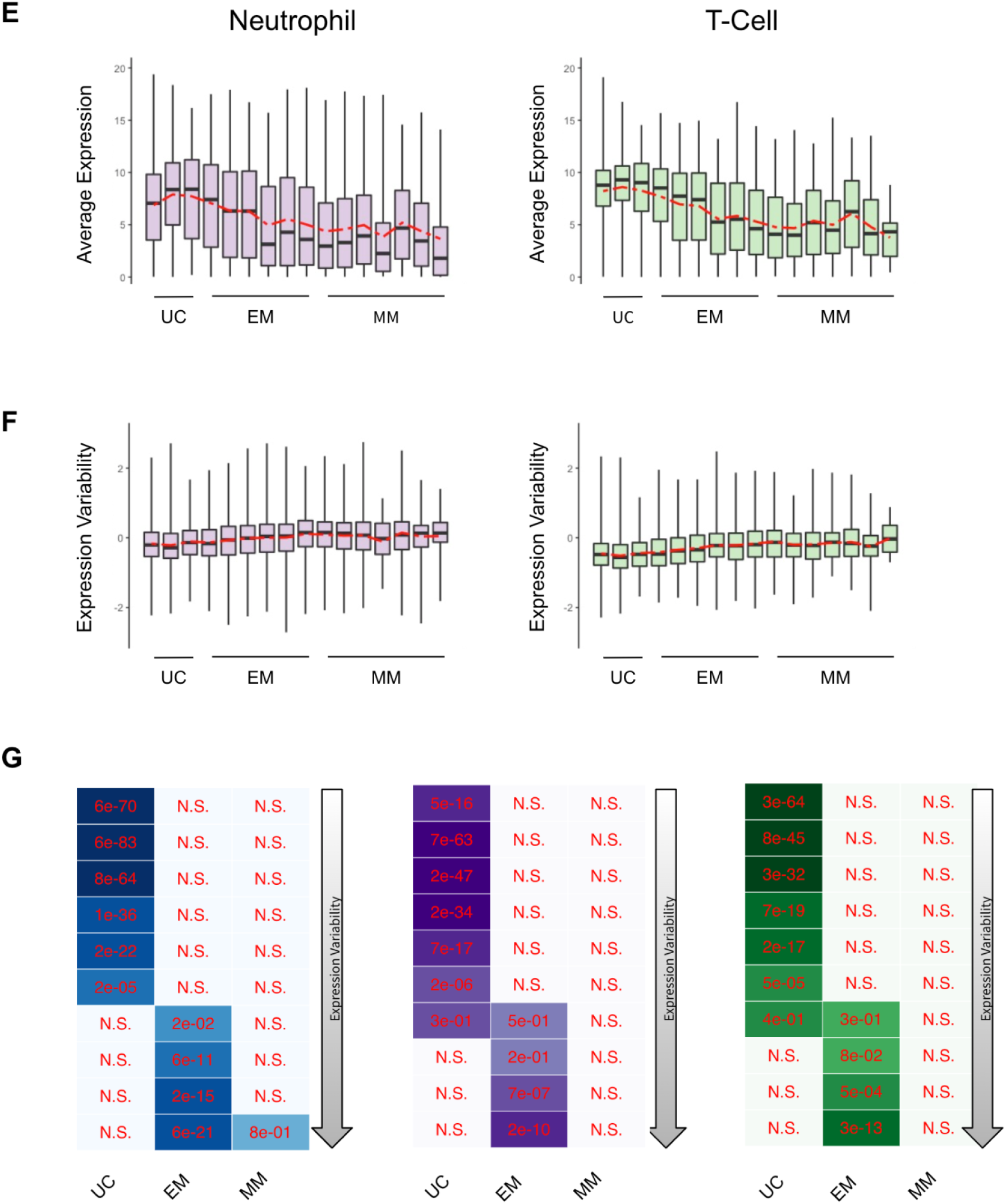
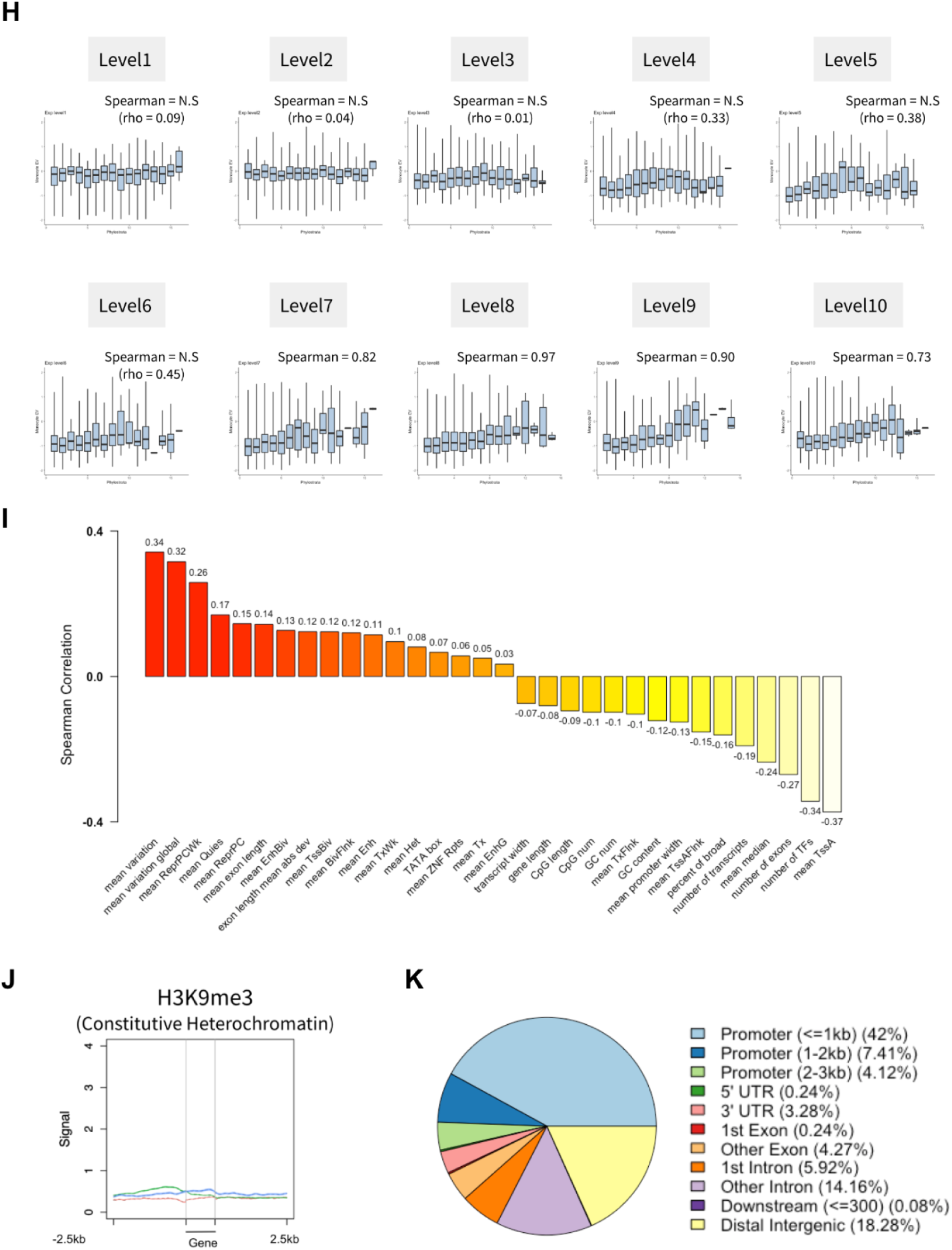
**A**) Number of genes in each phylostratum (left) and in each main age class (right). **B**) Gene Ontology enrichment of genes from each phylostratum. **C**) Proportion of housekeeping genes in each phylostratum (left) and in each main age class (right). **D**) Proportion of Leukemia proto-oncogenes in each phylostratum (left) and in each main age class (right). Dataset from Jang *et al*., 2015. **E**) Boxplot of average inter-individual gene expression levels in Neutrophil (left) and T-Cell (right) across 16 gene age classes (Spearman test in neutrophil, Rho = −0.88, *p*-value < 2.2e-16. Spearman test in t-cell, Rho −0.85, *p*-value = 1.023e-05). **F**) Boxplots of inter-individual expression variability in Neutrophil (left) and T-Cell (right) across 16 gene age classes (Spearman test in neutrophil, Rho = 0.74, *p*-value = 0.001703. Spearman test in t-cell, Rho = 0.84, *p*-value = 1.932e-05). **G**) Fisher’s Exact Tests of 10 EV decile levels and 3 main gene ages in Monocyte, Neutrophil and T-Cell, from left to right. Color shades show Jaccard index. This confirms the link between EV and gene evolutionary ages. **H**) Boxplots of inter-individual expression variability in Monocyte across the 16 phylostrata for 10 gene expression variability level classes. **I**) Spearman correlations of various DNA variability properties with 16 gene age classes. **J**) Average H3K9me3 profile in Monocyte around the 3 main gene age classes (Red Unicellular genes, Green Early metazoan genes, Blue Mammal Specific genes) (Sigalova et al., 2020). **K**) Genomic localisations of all detected CpG islands in monocytes.

**Supplementary Fig 2. (related to Fig. 2).**
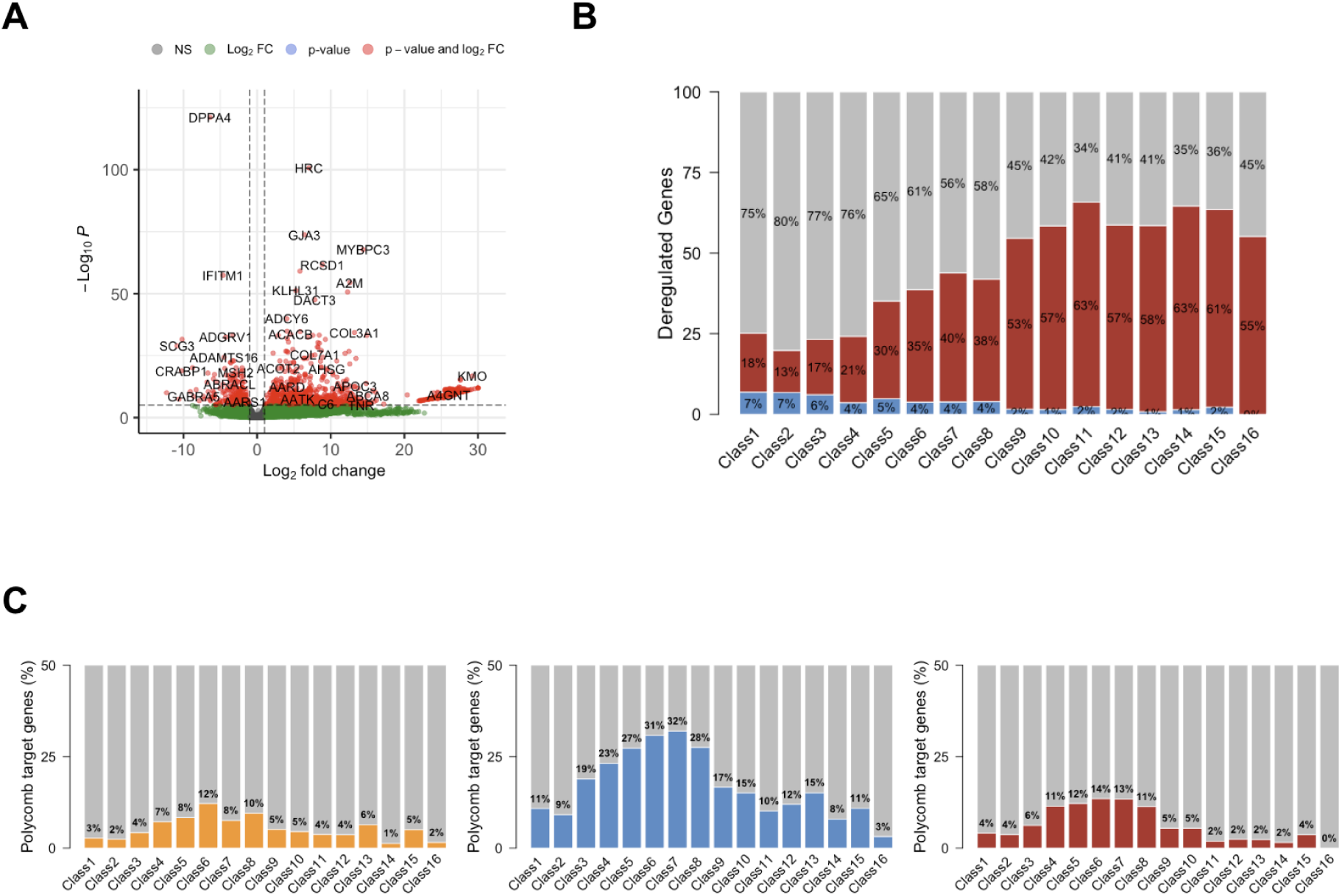
**A**) Volcano plot of differentially expressed genes in Cardiomyocyte compared to hESC. **B**) Proportion of Down (blue), Up (red) and non (grey) deregulated genes in Cardiomyocyte compared to hESC across the 16 gene age categories. **C**) Proportion of Polycomb target genes across the 16 gene age categories in hESC (left), B-cell (middle) and CLL (right).

**Supplementary Fig 3. (related to Fig. 3).**
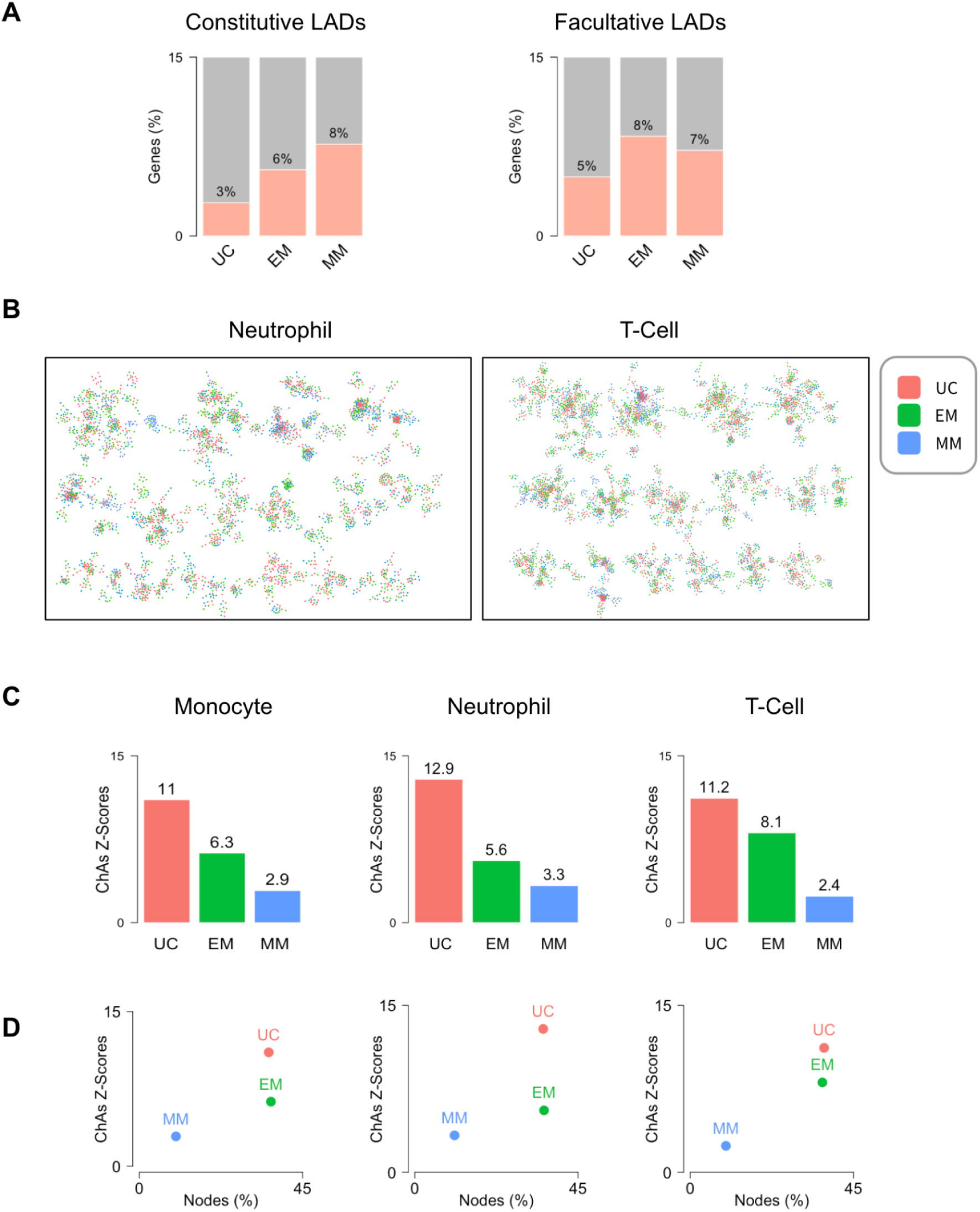
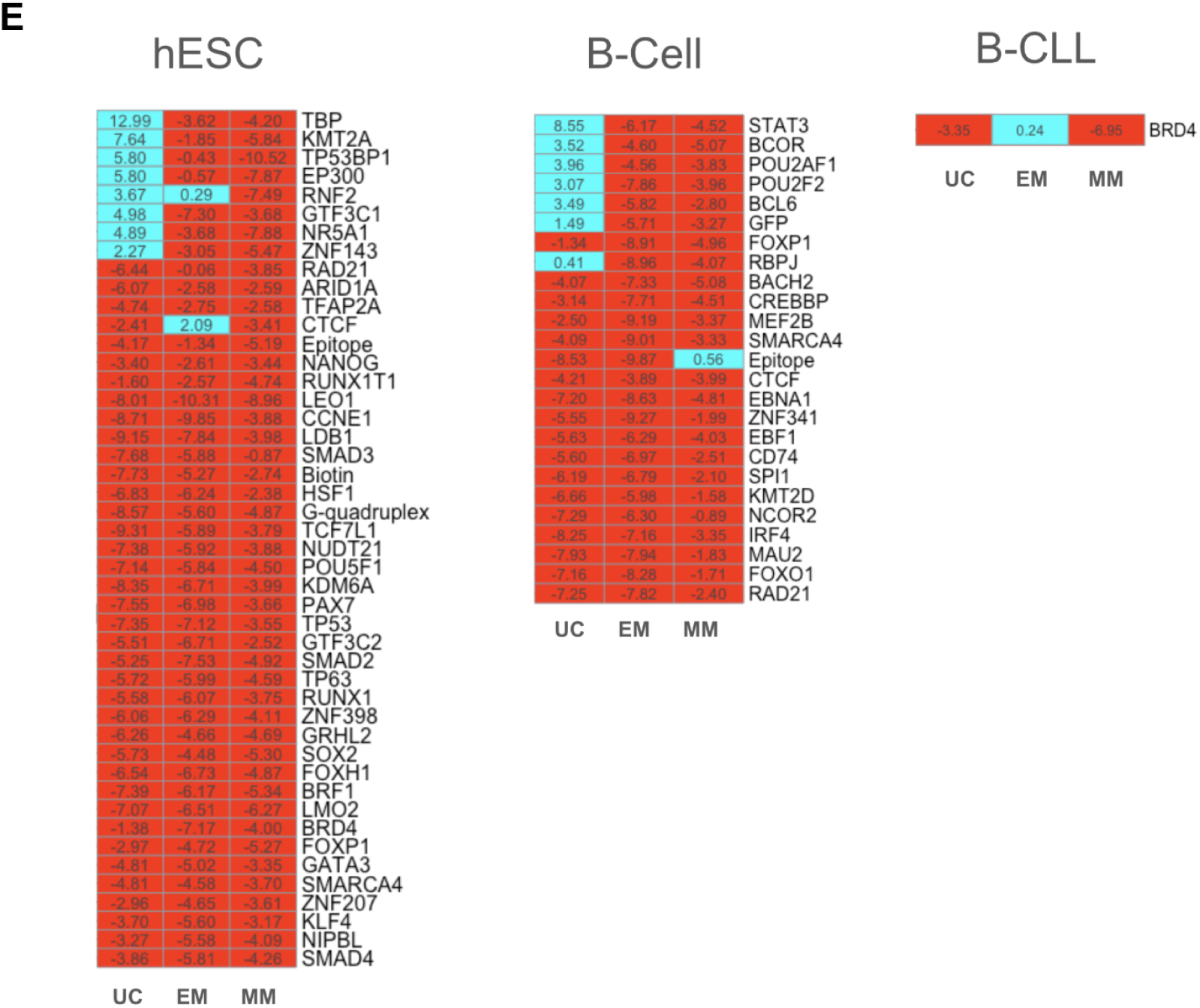
**A**) Overlaps with Lamina Associated Domains (LADs) annotated in multiple cell lines nd for genes belonging to the different age classes. **B**) Visualizations of the 3D promoter-promoter interaction networks in neutrophil (left) and T-cell (right) with node color representing gene age classes (red = UC, green = EM, blue = MM). **C**) ChAs z-scores compared amongst the 3 age classes in monocyte, neutrophil and T-cell promoter-promoter interaction networks. **D**) ChAs z-scores and related feature abundance of the 3 age classes in monocyte, neutrophil and T-cell promoter-promoter interaction networks. **E**) ΔChAs z-scores matrix of several transcription factors and chromatin features in hESC, B-Cell, and B-CLL. Colors represent ΔChAs > 0 in cyan and ΔChAs < 0 in red.

## Notes

### Competing Interest Statement

The authors have declared no competing interest.

### Summary of Updates

We modified the title, some figures, and some parts of the text.

